## Supplemental Files for "Costimulation of TLR8 responses by CXCL4 in Human Monocytes Mediated by TBK1-IRF5 Signaling and Epigenomic Remodeling"

**Supplementary figures: 11**

**Supplementary table 3**

**Supplementary table 4**

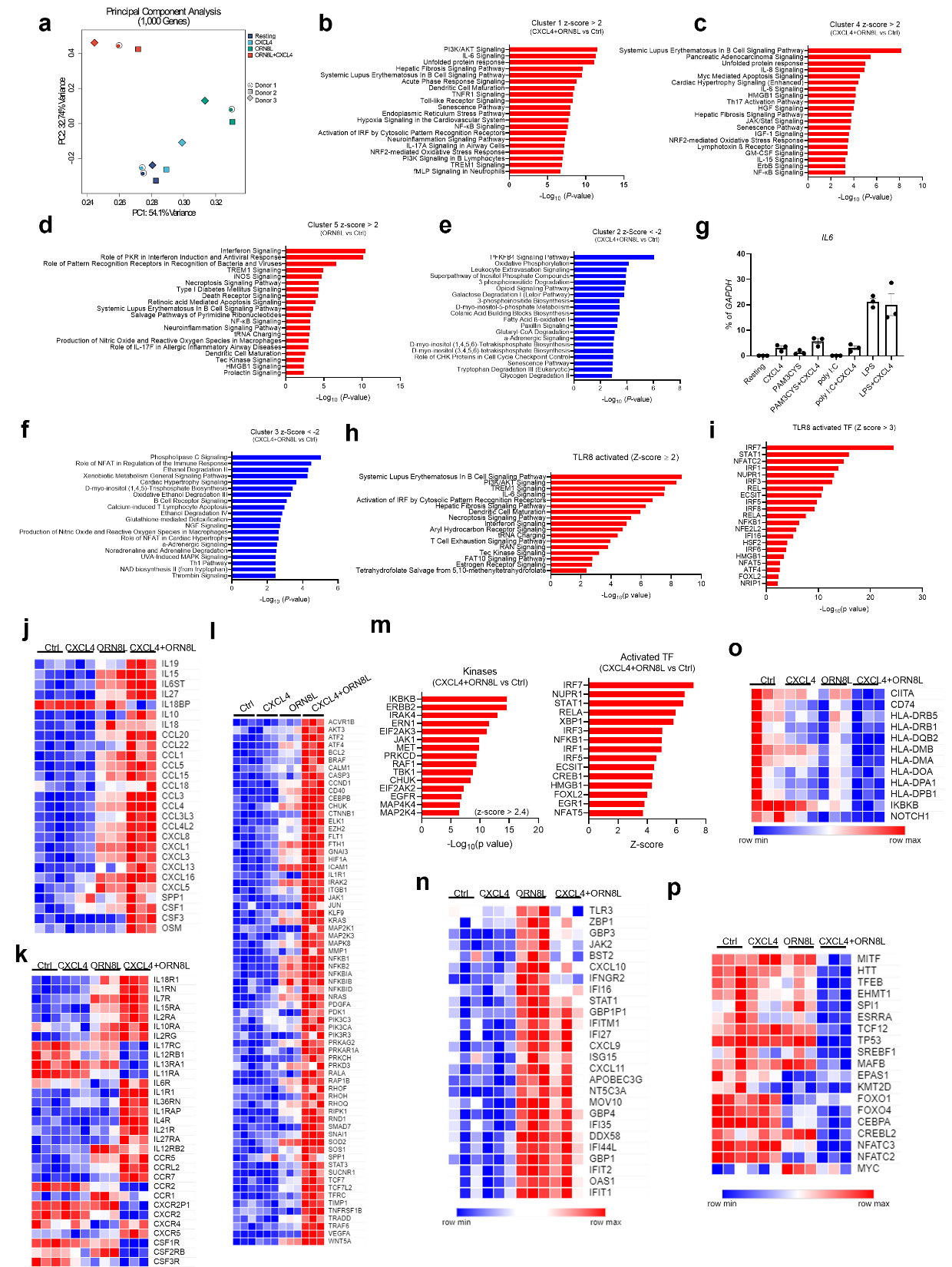

**Supplementary Figure 1.** (**a**) Principal Component Analysis (PCA) of RNAseq data. (**b** - **f, h, i, m**) IPA pathway analysis of gene clusters from **Figure 1b**. (**g**) mRNA of *IL6* was measured by quantitative PCR (qPCR) and normalized relative to *GAPDH* mRNA. n = 3. (**j** - **l, n, o**) Heatmaps showing expression of representative cytokine and chemokine genes (**j**) and their receptors (**k**) and fibrosis related genes (**l**), and ISGs (**n**), and antigen presentation related genes (**o**) and transcription factors important for anabolic metabolism and osteoclastogenesis (**p**) assessed in experimental condition as in **b** and presented (key) relative to the maximum. RNAseq was performed with three independent biological replicates.

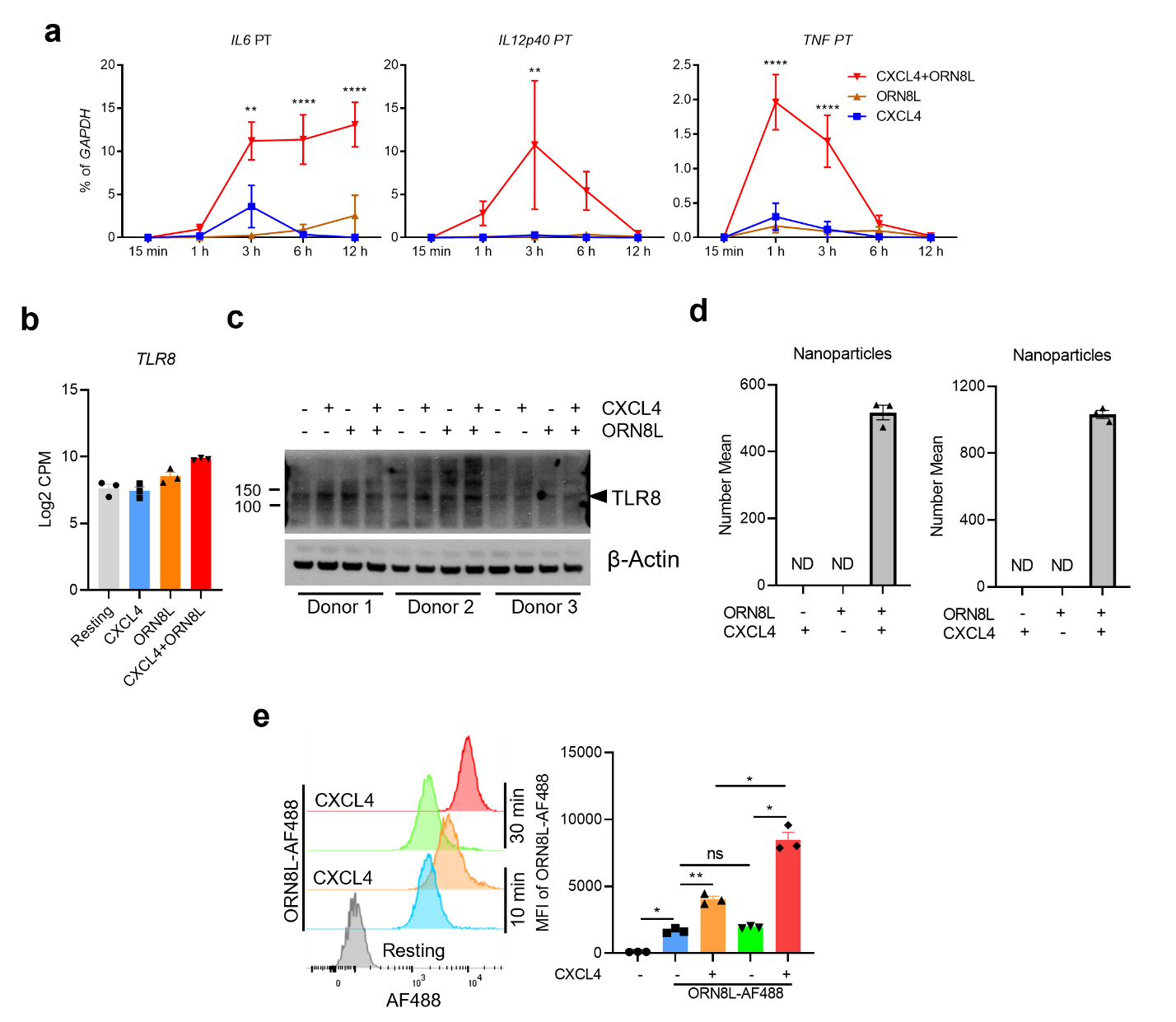

**Supplementary Figure 2.** (**a**) Primary transcripts (PT) of *IL6*, *IL12B* and *TNF* were measured by quantitative PCR (qPCR) and normalized relative to *GAPDH* mRNA. (**b**) Log2 CPM of TLR8 from RNAseq data in human monocytes 6 hr after CXCL4 and/or ORN8L stimulation. (**c**) Immunoblots of TLR8 using whole cell lysates from the indicated conditions 6 hr after stimulation (n = 3). (**d**) Nanoparticle formation measured using dynamic light scattering. The number mean is from one sample measured in triplicate. ND = not detected. 2 independent experiments are shown to supplement the third experiment shown in Fig. 1f. (**e**) Flow cytometric analysis of the internalization of ORN8L-AF488 after the indicated times of incubation in the absence or presence of CXCL4 in human monocytes. Left panel, representative FACS plot; right panel, cumulative data. Data is depicted as mean ± SEM; ****p ≤ 0.0001; ***p ≤ 0.001; **p ≤ 0.01; *p ≤ 0.05 by two-way ANOVA (**a**) or one-way ANOVA (**e**).

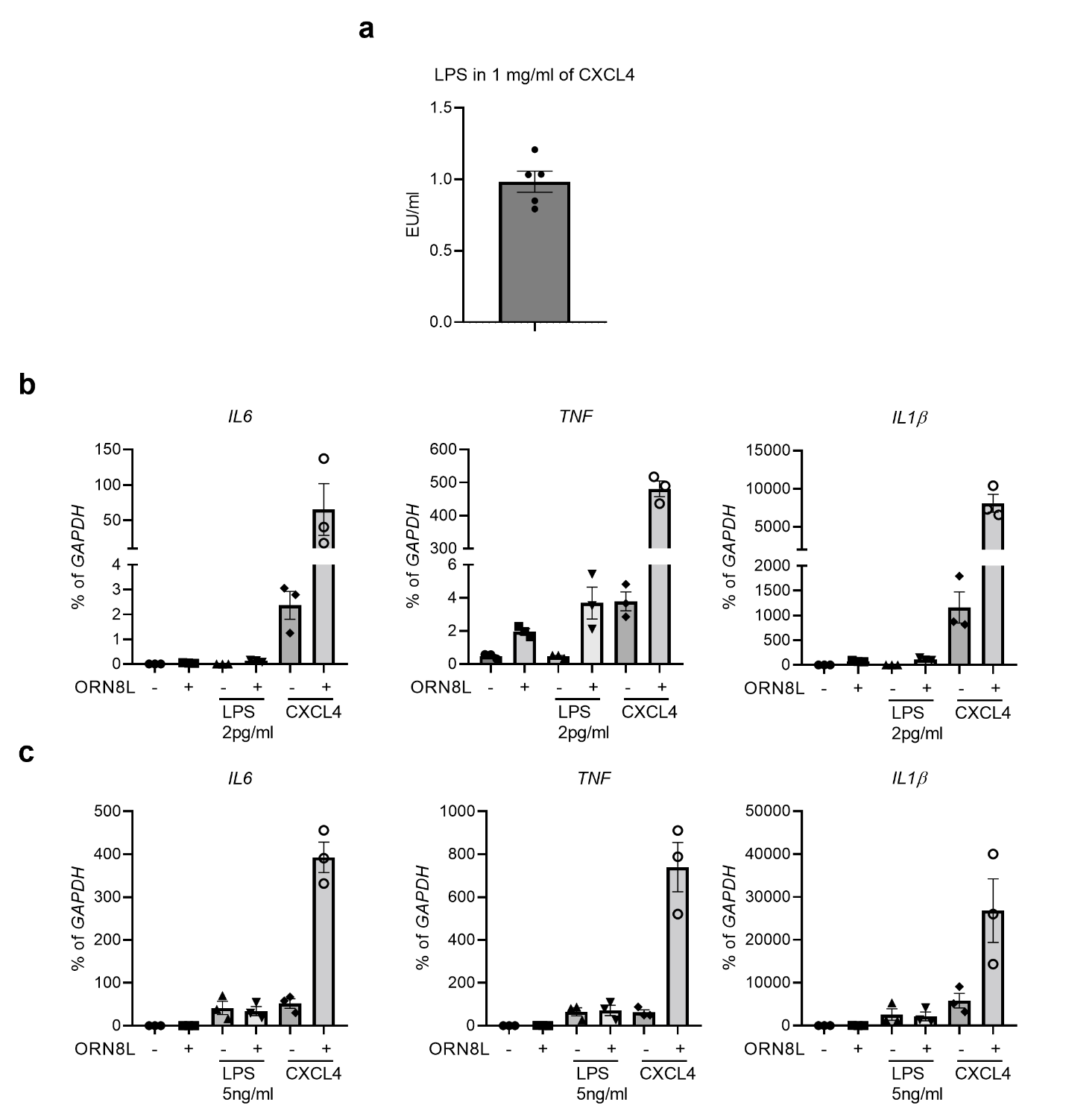

**Supplementary Figure 3.** (**a**) Endotoxin assay of CXCL4 stock solution. (**b** and **c**) qPCR analysis of mRNA amounts of *IL6*, *TNF* and *IL1β* normalized relative to *GAPDH* mRNA in cells stimulated with 2 pg/ml LPS (**b**) or 5 ng/ml LPS (**c**) in the presence/absence of ORN8L, CXCL4 serves as a positive control.

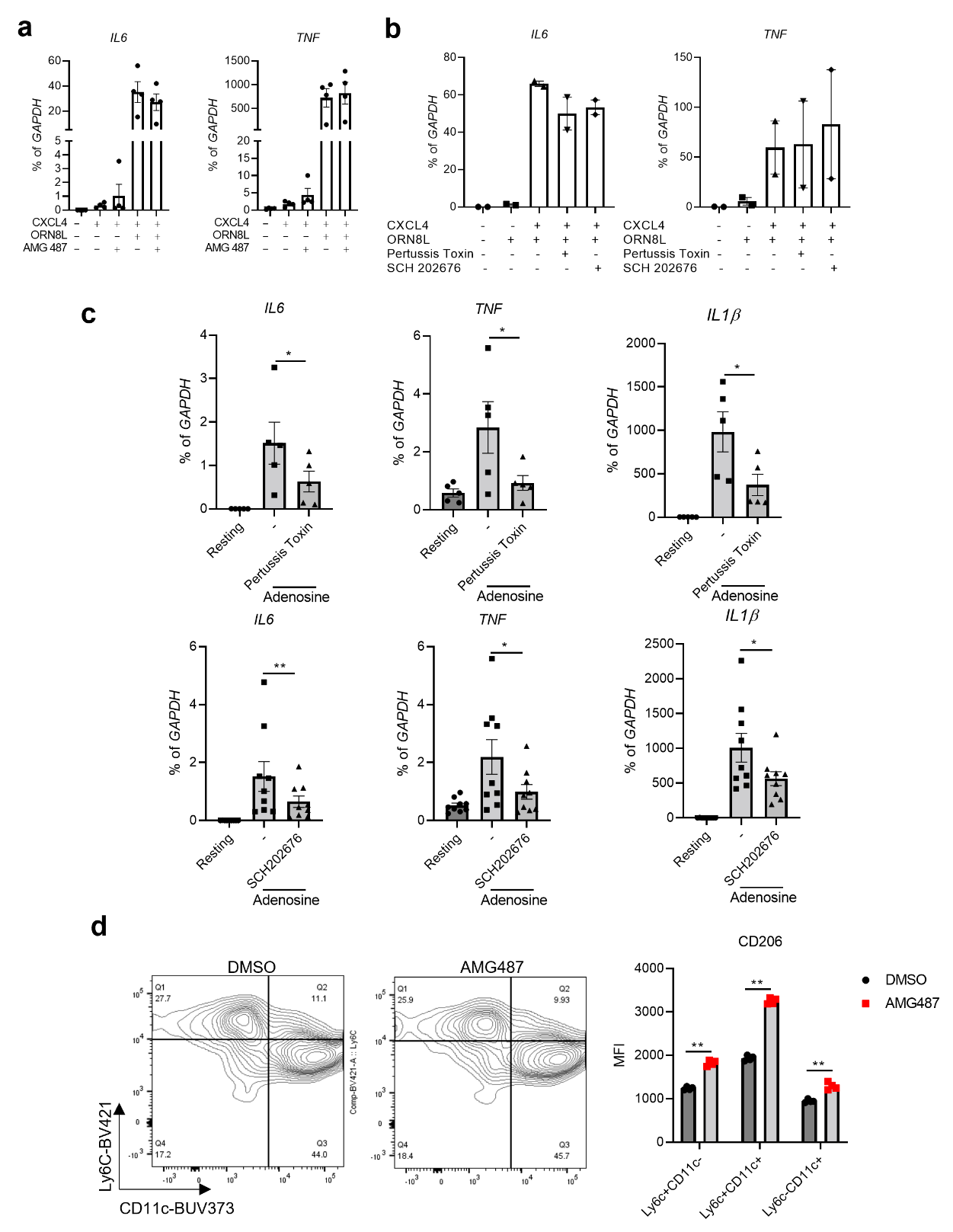

**Supplementary Figure 4.** (**a**) mRNA of *IL6* and *TNF* was measured by qPCR and normalized relative to *GAPDH* mRNA in cells stimulated with CXCL4 and/or ORN8L with/without CXCR3 inhibitor AMG487 (5 μM) for 3 hr (n = 4). (**b** and **c**) mRNA of *IL6* and *TNF* was measured by qPCR and normalized relative to *GAPDH* mRNA in cells stimulated with CXCL4 and/or ORN8L (**b**, n = 2) or adenosine (1mM) (**c**, n = 5-9) with/without G-coupled receptor inhibitors Pertussis toxin (10 μM) or SCH202675 (10 μM) for 6 h. (**d**) FACS to analyze CD206 levels in BMDCs with/without CXCR3 inhibitor AMG487 (5 μM) for 24 hr (n = 4). Data depict mean ± SEM; **p ≤ 0.01; *p ≤ 0.05 by paired t test (**c**) or two-way ANOVA (**d**). (**c**) and (**d**) provide positive controls for efficacy of the inhibitors used.

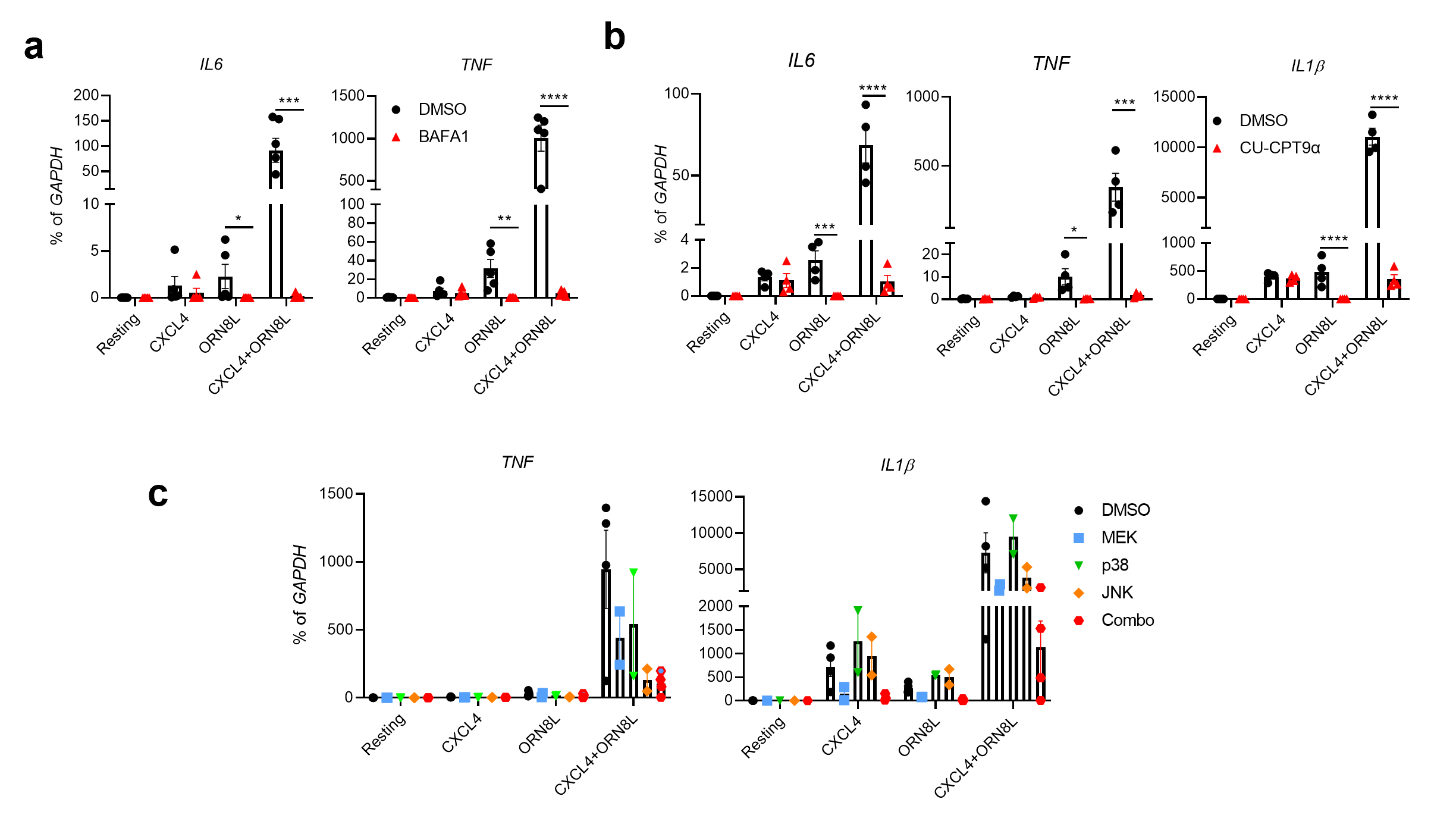

**Supplementary Figure 5.** (**a** and **b**) mRNA of *IL6*, *TNF* and *IL1β* was measured by qPCR and normalized relative to *GAPDH* mRNA in cells stimulated with CXCL4 and/or ORN8L with/without Bafilomycin A1 (BAFA1) (1 μM), TLR8 inhibitor CU-CPT9α (1 μM) for 3 h, respectively. (**c**) qPCR analysis of mRNA amounts of *TNF* and *IL1β* normalized relative to *GAPDH* mRNA in cells stimulated with CXCL4 and/or ORN8L after with treatment the MAPK inhibitors SB 202190 (p38), JNK inhibitor II and U0126 (MEK1/2) used at 10 μM, respectively. The data are related to and use some of the same samples as Fig. 2g. Data is depicted as mean ± SEM of 2 - 4 healthy donors; ****p ≤ 0.0001; ***p ≤ 0.001; **p ≤ 0.01; *p ≤ 0.05 by two-way ANOVA.

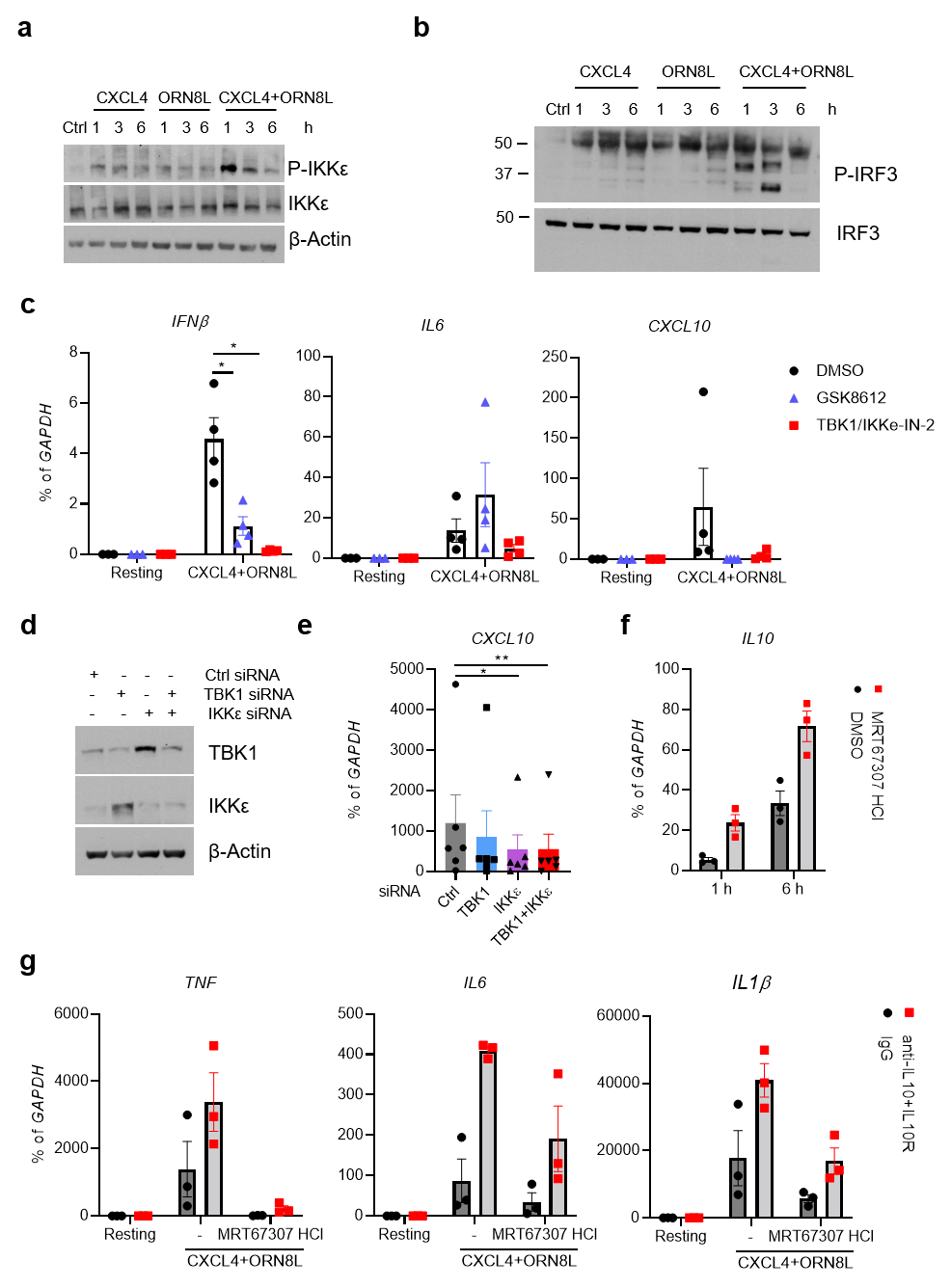

**Supplementary Figure 6.** (**a** and **b**) Immunoblots of phospho-IKKε, total IKKε (**a**) and phospho-IRF3 and total IRF3 (**b**) using whole cell lysates from a time course with the indicated conditions. (**c**) mRNA of indicated genes was measured by qPCR and normalized relative to *GAPDH* mRNA after blockade of TBK1/IKKε activation by 1 μM of TBK1/IKKε-IN-2 or 50 μM of GSK8612 (n = 4 independent experiments) for 3 h. (**d**) Immunoblot of TBK1 and IKKε with whole cell lysates from monocytes nucleofected with control or TBK1- and/or IKK-specific siRNAs. (n = 3) (**e**) mRNA of *CXCL10* was measured by qPCR and normalized relative to *GAPDH* mRNA using monocytes nucleofected with control or TBK1- and/or IKK-specific siRNAs. (**f**) mRNA level of *IL10* measured by qPCR and normalized relative to *GAPDH* mRNA (n = 3 independent experiments). (**g**) mRNA of indicated genes was measured by qPCR and normalized relative to *GAPDH* mRNA after blockade of IL-10 and IL10R with/without TBK1 inhibitor MRT67307 HCl (n = 3 independent experiments). Data in (**a** and **b**) are representative of 3 experiments. **p ≤ 0.01; *p ≤ 0.05 by two-way ANOVA (**c**) or by Friedman test (**e**).

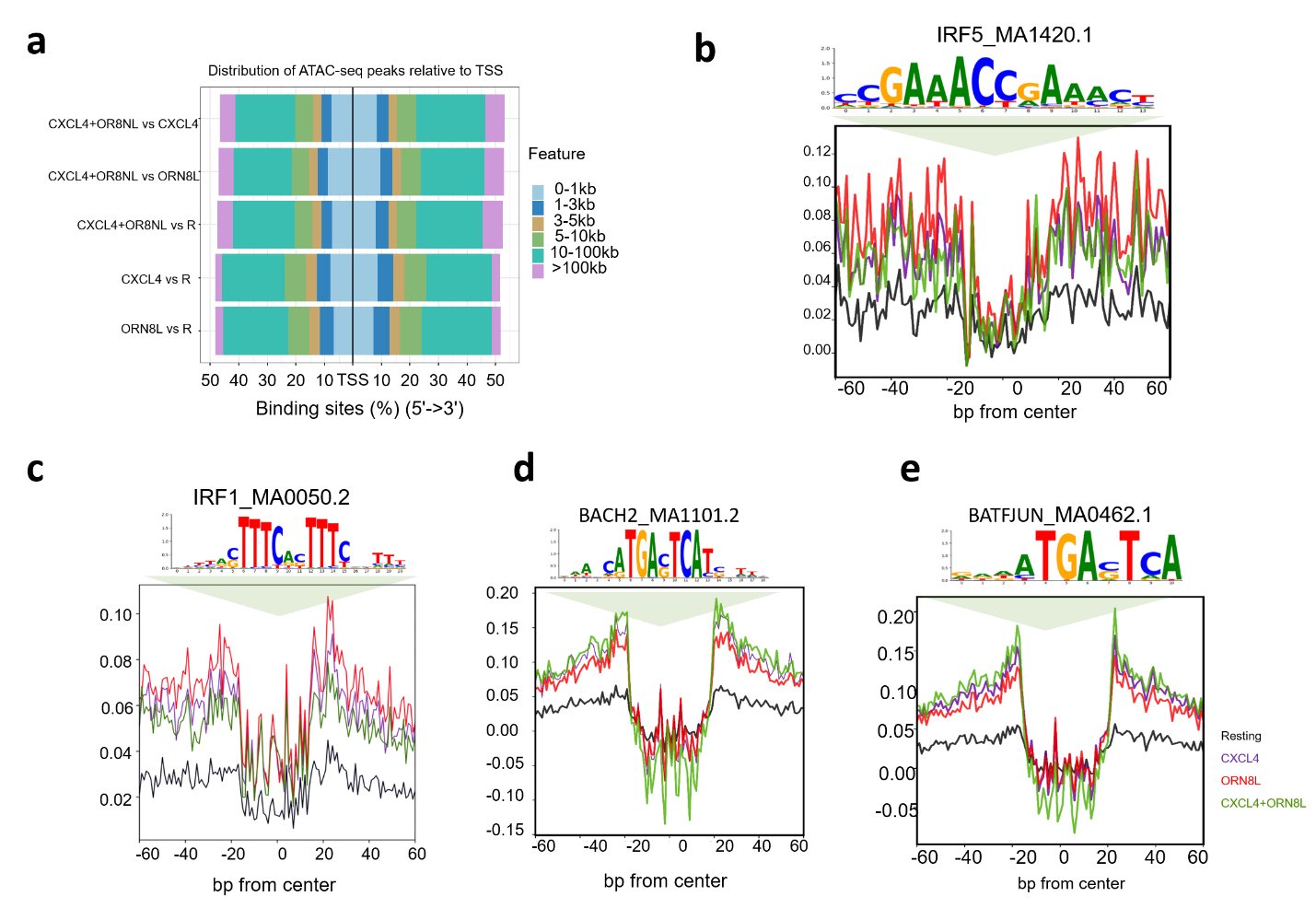

**Supplementary Figure 7.** (**a**) Pie chart representation of the relative distribution of ATAC-seq peak coordinates relative to the position of transcription start sites (TSS) for differential open chromatin regions identified by comparison of respective treatments versus the resting condition. (**b** - **e**) Visualization of dignificant footprints for all accessible sites for the following significant motifs IRF1_MA0050.2, IRF5_MA1420.1, BACH2_MA1101.2, BATFJUN_MA0462.1 that were identified by BINDetect using TOBIAS.

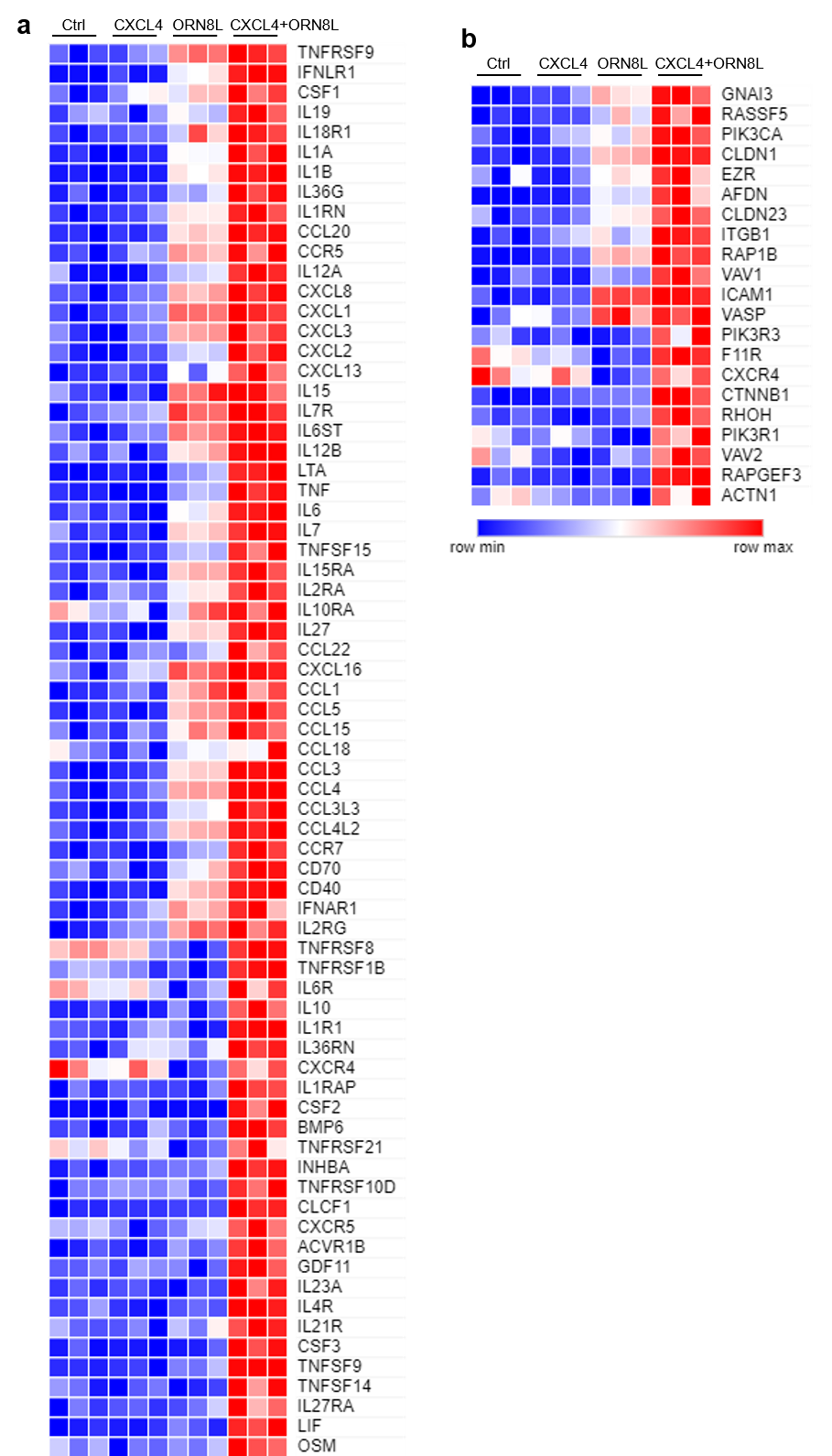

**Supplementary Figure 8.** (**a**) Heatmap showing representative genes associated with C1 peaks in the cytokine-cytokine receptor pathway. (**b**) Heatmap showing representative genes associated with C1 peaks in cell adhesion and migration pathways.

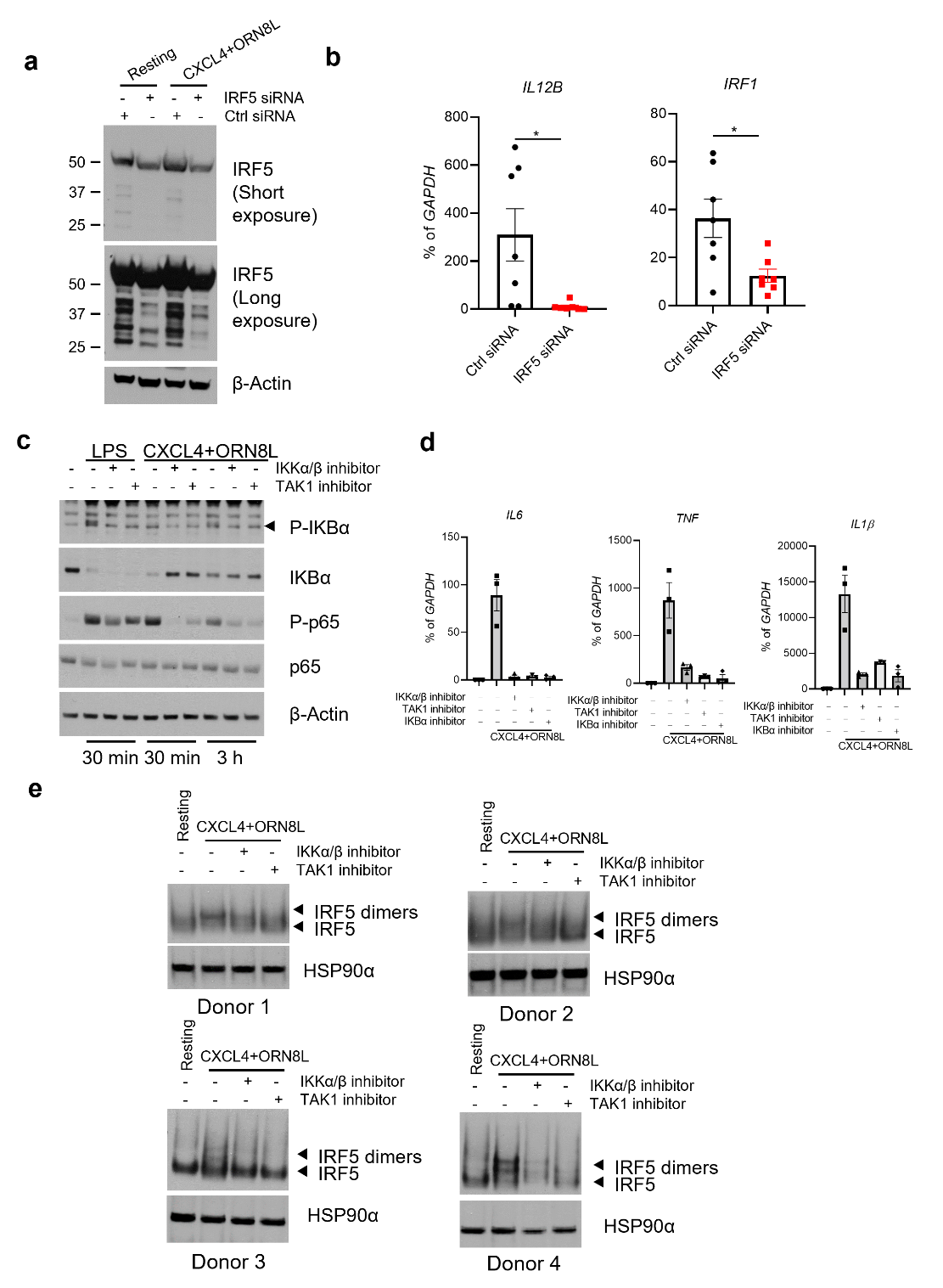

**Supplementary Figure 9.** (**a**) Immunoblot of IRF5 using whole cell lysates 3 days after nucleofection of monocytes with control or IRF5-specific siRNAs. (**b**) mRNA of indicated genes was measured by qPCR and normalized relative to *GAPDH* mRNA after knockdown of IRF5 by siRNA for 3 days (n = 7 independent experiments). Data are representative of 3 independent experiments (**a**) or depict mean ± SEM (**b**). *p ≤ 0.05 by Wilcoxon signed-rank test. (**c**) Immunoblots of phospho-IKBα, total IKBα, and phospho-p65 and total p65 using whole cell lysates from the indicated conditions. (**d**) mRNA of *IL6*, *TNF* and *IL1β* was measured by qPCR and normalized relative to *GAPDH* mRNA in cells stimulated with CXCL4 and/or ORN8L and IKKα/β inhibitor BMS-345541 (10 μM), TAK1 inhibitor Takinib (10 μM), and IKBα inhibitor Bay 11-7085 (10 μM), respectively. (**e**) Immunoblot of IRF5 using whole cell lysates of indicated conditions run on nondenaturing gels. HSP90α serves as loading control. Four independent experiments are shown.

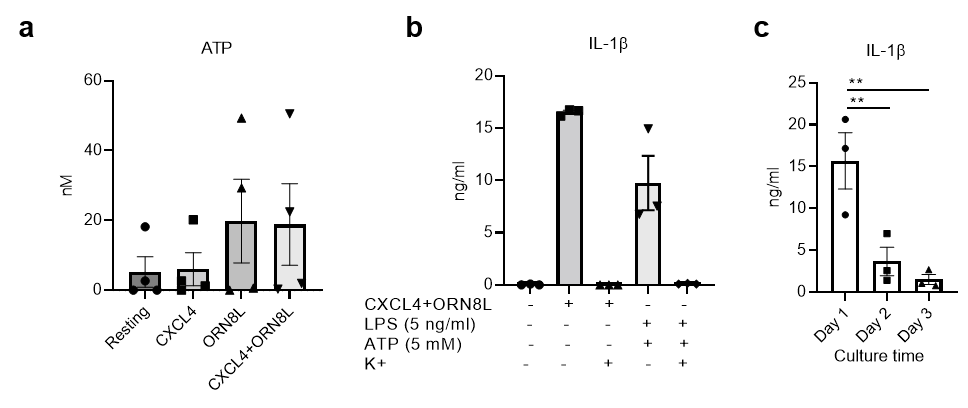

**Supplementary Figure 10.** (**a**) ATP concentration in indicated cell culture medium detected using ATP Determination Kit. (**b**) 130 mM extracellular concentration of K+ suppressed (CXCL4 + ORN8L)-induced IL-1β secretion. (**c**) ELISA of IL-1β protein in culture supernatants of monocytes cultured in presence of M-CSF for 1-3 days prior to CXCL4 and TLR8 costimulation for 6h. Data is depicted as mean ± SEM of 3 healthy donors; **p ≤ 0.01 by one-way ANOVA.

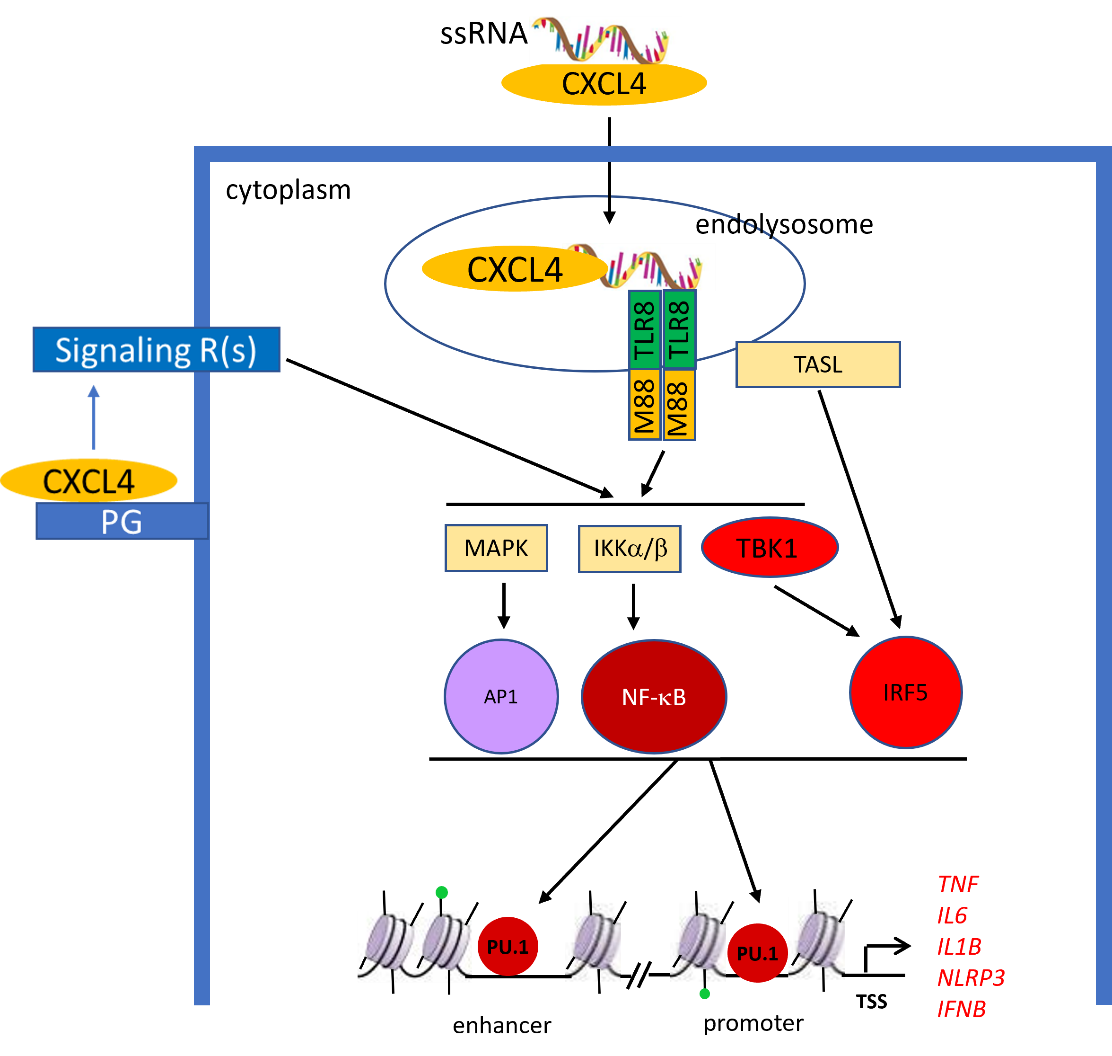

**Supplementary Figure 11.** A schematic model linking CXCL4 and TLR8 signaling with chromatin remodeling and de novo enhancers associated with inflammatory genes.

**Supplementary table 3: Primers and siRNAs.**

| **Primers for qPCR Source** | | |
| --- | --- | --- |
| Human *GAPDH* F: ATCAAGAAGGTGGTGAAGCA; R: GTCGCTGTTGAAGTCAGAGGA | This paper |  |
| Human *IL6* F: TAATGGGCATTCCTTCTTCT; R: TGTCCTAACGCTCATACTTTT | This paper |  |
| Human *IL12B* F: GGGCACAGATGCCCATTCGCT; R: GGGCACAGATGCCCATTCGCT | This paper |  |
| Human *TNF* F: AATAGGCTGTTCCCATGTAGC; R: AGAGGCTCAGCAATGAGTGA | This paper |  |
| Human *IL6* primary transcript (PT) F: TTGCAAGGAAGGTTTTTGGAG  R: CTTGGGTTCAGTTCCAAGCTC | This paper |  |
| Human *IL12B* primary transcript (PT) F: AAGCACTTTGGAGGAAGCATAG; R: TGAACCCAAGTCAATGTGAGTC | This paper |  |
| Human *TNF* primary transcript (PT) F: TAAGGGTGACTCCCTCGATGT; R: CCAAACCCAAACCCAGAATTA | This paper |  |
| Human *IL1β* F: TTCGACACATGGGATAACGAGG; R: TTTTTGCTGTGAGTCCCGGAG | This paper |  |
| Human *CXCL10* F: TTAATCTTGTCTCTGGGCTTGG; R: GTTGGGGAATGAGGTTAGGG | This paper |  |
| Human *IRF1* F: GCACTAAGCGAAAATTGCA; R: GGGAGTTTCCTTCACATTCA | This paper |  |
| Human *NLRP3* F: GATCTTCGCTGCGATCAACAG; R: CGTGCATTATCTGAACCCCAC | This paper |  |
| Mouse *Gapdh* F: ATCAAGAAGGTGGTGAAGCA; R: AGACAACCTGGTCCTCAGTGT | This paper |  |
| Mouse *Il6* F: TGGCTAAGGACCAAGACCATCCAA; R: AACGCACTAGGTTTGCCGAGTAGA | This paper |  |
| Mouse *Tnf* F: CCCTCACACTCAGATCATCTTCT; R: GCTACGACGTGGGCTACAG | This paper |  |
| Mouse *Il1β* F: AGCTTCCTTGTGCAAGTGTCT; R: GACAGCCCAGGTCAAAGGTT | This paper |  |
| **FAIRE Primers for qPCR** |  |  |
| Human *IL6* (promoter) F: ACCCTCACCCTCCAACAAAG; R:  GCAGAATGAGCCTCAGACATC | This paper |  |
| Human *TNF* (promoter) F: GCCCCAGGGACATATAAAGG; R: GCCCCAGGGACATATAAAGG | This paper |  |
| Human *IL12B* (promoter) F: CCCAGAAGGTTTTGAGAGTTGT; R: GATGTTGTTTCTTCTGCTGCTG | This paper |  |
| **Oligonucleotides Source CAS#** | | |
| siGENOME Human TBK1 siRNA | Horizon | M-003788-02-0005 |
| siGENOME Human IKBKE siRNA | Horizon | M-003723-02-0005 |
| siGENOME Human IRF5 siRNA | Horizon | M-011706-00-0005 |
| ON-TARGETplus Human MYD88 siRNA | Horizon | L-004769-00-0005 |
| ON-TARGETplus Human TICAM1 siRNA | Horizon | L-012833-00-0005 |

**Supplementary table 4: Antibodies and Others.**

| **Antibodies Source CAS#** | | |
| --- | --- | --- |
| IkBa | Cell Signaling Technology | 9242s |
| p-p38 | Cell Signaling Technology | 9215S |
| anit-p38 | Cell Signaling Technology | 9212S |
| Phospho-p44/42 MAP Kinase (ERK1/2) | Cell Signaling Technology | 9101S |
| ERK1/2 | Cell Signaling Technology | 9102S |
| TLR8 | Thermofisher Scientific | PA5-80137 |
| TBK1/NAK (E8I3G) | Cell Signaling Technology | 38066S |
| Phospho-TBK1/NAK (Ser172) (D52C2) | Cell Signaling Technology | 5483T |
| Phospho-IKKε (Ser172) (D1B7) | Cell Signaling Technology | 8766S |
| IKKε | Cell Signaling Technology | 2690T |
| NF-κB p65 (D14E12) | Cell Signaling Technology | 8242S |
| Phospho-NF-κB p65 (Ser536) (93H1) | Cell Signaling Technology | 3033S |
| p-IKBα | Cell Signaling Technology | 9246S |
| IKBα | Cell Signaling Technology | 9242S |
| Phospho-IRF-3 (Ser396) | Cell Signaling Technology | 29047S |
| IRF3 | Cell Signaling Technology | 11904T |
| IRF5 Polyclonal Antibody | Invitrogen | PA5-19504 |
| Phospho-CYLD (Ser418) Antibody | Cell Signaling Technology | 4500T |
| β-Actin (D6A8) Rabbit mAb | Cell Signaling Technology | 8457 |
| NLRP3 (D4D8T) Rabbit mAb | Cell Signaling Technology | 15101 |
| Human IL-1 beta /IL-1F2 Antibody | R&D Systems | MAB601R-100 |
| Caspase-1 (D7F10) Rabbit mAb | Cell Signaling Technology | 3866 |
| Cleaved Gasdermin D (Asp275) (E7H9G) | Cell Signaling Technology | 36425 |
| AIM2 (D5X7K) Rabbit mAb | Cell Signaling Technology | 12948 |
| Human IL-1 beta /IL-1F2 Biotinylated Antibody | R&D Systems | BAF201 |
| Phospho-IRF-3 (Ser386) (E7J8G) XP® Rabbit mAb (Alexa Fluor® 488 Conjugate) | Cell Signaling Technology | 73981 |
| Human/Primate IL-6 Antibody | R&D Systems | MAB206-SP |
| Human/Primate IL-6 Biotinylated Antibody | R&D Systems | BAF206 |
| Human TNF-alpha Antibody | R&D Systems | MAB610-SP |
| Human TNF-alpha Biotinylated Antibody | R&D Systems | BAF210 |
| **TLR ligands, inhibitors and recombinant proteins** | | |
| Recombinant Human IL-1 beta/IL-1F2 Protein | R&D Systems | 201-LB-005 |
| LPS | Invivogen | TLRL-3pelps |
| PAM3CYS | Invivogen | TLRL-PMS |
| Poly I:C | Invivogen | TLRL-PIC |
| ORN8L | Chemgenes Corporation | (Lan et al., 2007) |
| ORN8L-AF488 | Chemgenes Corporation |  |
| Recombination Human CXCL4 | PEPROTECH | 300-16 |
| PF-4 (CXCL4) human | Sigma-Aldrich | SRP3142 |
| Recombinant Human IL-6 | Peprotech | 200-06 |
| Recombinant Human TNF-alpha | Peprotech | 300-01A |
| Pertussis Toxin | R&D Systems | 3097/50U |
| SCH 202676 hydrobromide | R&D Systems | 1400/10 |
| BMS-345541-IKKα/β inhibitor | Selleckchem | S8044 |
| Bay 11-7085-IKBα inhibitor | Selleckchem | S7352 |
| Takinib | Selleckchem | S8663-5mg |
| NLRP3 Inhibitor, MCC950 | Sigma Aldrich | 5381200001 |
| Ac-YVAD-cmk, CASP1 inhibitor | Sigma Aldrich | SML0429 |
| SB202190, Hydrochloride, p38 inhibitor | Calbiochem | 559393 |
| Bafilomycin A1 | MCE | HY-100558 |
| MRT67307 HCl (dual IKKϵ and TBK1 inhibitor) | Selleckchem | S7948 |
| TBK1/IKKε-IN-2 | MCE | HY-12453 |
| GSK8612-TBK1 inhibitor | Selleckchem | S8872 |
| AMG 487 CXCR3 antagonist | TOCRIS | 4487 |
| JNK Inhibitor II | Sigma Aldrich | 420119 |
| U0126 MEK1/2 inhibitor | Sigma Aldrich | 662005 |
